# FraCeMM - A Framework for Cell–Matrix Mechanotransduction

**DOI:** 10.64898/2026.03.16.712065

**Authors:** Igor Nelson

## Abstract

Cells sense and respond to the mechanical properties of their environment, yet the minimal physical principles sufficient to reproduce mechanotransduction and durotaxis remain debated. This work introduces FraCeMM, a physics-first mechanochemical simulation framework coupling stochastic ligand–integrin–talin binding to a deformable soft-body cell model on an elastic substrate. Without imposed polarity, directional cues, or migration rules, the model reproduces hallmark mechanobiological behaviors including stiffness-dependent spreading, traction reinforcement, focal adhesion asymmetry, and directed durotaxis. A finite pool of adhesion molecules, mechanically coupled through elastic linkages, drives emergent force asymmetry and polarization via self-consistent feedback between stochastic binding, molecular availability, and substrate stiffness. Despite minimal assumptions and a coarse-grained molecular representation, resulting traction forces, adhesion loads, and migration speeds fall within experimentally reported ranges. These results support the view that local force balance, limited adhesion resources, and mechanically binding are sufficient to generate adaptive mechanosensing and directed migration, establishing a transparent and extensible foundation for computational mechanobiology.

## 1. Introduction

Focal adhesions (FAs) are specialized, dynamic macromolecular assemblies that form at the interface between a cell and its extracellular matrix (ECM) Hynes (2002); Geiger and Yamada (2011). They function simultaneously as mechanical linkages and signaling hubs, anchoring the actin cytoskeleton to the ECM while transducing mechanical cues into biochemical signals that regulate essential cellular processes Sun et al. (2016).

Structurally, FAs comprise integrins, talin, vinculin, actin filaments, and numerous adapter and signaling proteins that together form a highly organized nanoscopic architecture bridging the intracellular and extracellular environments Geiger and Yamada (2011).

Through these structures, cells adhere, migrate, and sense their surroundings. FAs transmit cytoskeletal tension to the ECM, enabling cells to exert traction forces required for motility, polarization, and mechanosensing Sun et al. (2016). They also regulate intracellular signaling pathways governing proliferation, differentiation, and survival Hynes (2002). Importantly, focal adhesions are not static entities; they continuously assemble, disassemble, and remodel in response to mechanical stress, substrate stiffness, and biochemical gradients Geiger and Yamada (2011). This dynamic turnover permits adaptation to diverse microenvironments and underlies processes such as *durotaxis*, the directed migration toward stiffer regions of the ECM Plotnikov and Waterman (2013).

Dysregulation of FA dynamics is implicated in numerous pathological conditions. In cancer, aberrant adhesion signaling contributes to uncontrolled proliferation and metastasis; in cardiovascular and developmental disorders, altered FA composition and mechanics compromise tissue integrity and function Hynes (2002). Understanding the biophysical principles that govern FA formation and stability therefore remains central to elucidating how mechanical and biochemical signals integrate to shape cellular behavior Sun et al. (2016).

### 1.1. Modeling Focal Adhesion Dynamics

The interplay between molecular binding, force generation, and mechanical feedback within focal adhesions presents a major challenge for quantitative biology (Iskratsch et al. 2014; Schwarz and Safran 2013). Modeling efforts have sought to bridge these processes using approaches that range from continuum mechanical descriptions to stochastic molecular simulations (Bell 1978; Erdmann and Schwarz 2019). Such models aim to reproduce experimentally observed behaviors, including cell spreading, elongation, and durotaxis, by capturing the coupled evolution of biochemical signaling and physical deformation (Plotnikov and Waterman 2013; Kim et al. 2021).

A notable contribution was presented, who combined a Cellular Potts model with an infinite element framework to examine how ECM stiffness modulates cell traction forces and FA dynamics (Rens and Merks 2020). Their results demonstrated that ECM stress amplifies traction forces, promoting cell elongation rather than adhesion stabilization. On compliant substrates, the buildup of force was insufficient to stabilize adhesions, leading to rounded cell morphologies. Conversely, intermediate stiffness facilitated durotaxis driven by asymmetric adhesion lifetimes across the substrate. These findings suggested that cells sense matrix stiffness through the rate of force development: as myosin motors generate contractile tension, the rate of tension increase, which depends on substrate stiffness, dictates FA assembly and disassembly kinetics (Plotnikov and Waterman 2013). This feedback mechanism explains phenomena such as enhanced cell spreading on stiff substrates, elongation on intermediate stiffness, and migration toward stiffer regions. At very high stiffness values, however, FA growth saturated, imposing an upper limit on migration velocity and revealing a fundamental mechanical constraint in the cell–ECM feedback loop (Rens and Merks 2020).

Complementary to this mechanical framework, Blücher *et al*. (2014) introduced a stochastic model describing the molecular-scale events underlying FA initiation (Blucher et al. 2014*a*). Using Gillespie’s algorithm, they simulated the probabilistic dynamics of integrin–ligand–talin (LIT) interactions during early adhesion formation. In this approach, the system state comprises molecular species connected through a stoichiometry matrix, with reaction propensities determining the probability of each event (Gillespie 1977). The stochastic simulation proceeds by calculating propensities, drawing random numbers to determine which reaction occurs and when, and iterating until the desired time is reached. This formalism captures intrinsic fluctuations in binding kinetics, providing insight into how molecular noise influences adhesion stability and maturation (Erdmann and Schwarz 2019). By linking random microscopic events to macroscopic adhesion behavior, such models offer a bridge between stochastic molecular interactions and emergent cell-scale mechanics.

### 1.2. Integrating Multiscale Approaches

Recent efforts have sought to unify mechanical and stochastic perspectives into multiscale frameworks capable of describing the full spectrum of FA behavior. Examples include hybrid models that couple continuum mechanics of the cytoskeleton and ECM with stochastic adhesion kinetics (Walcott and Sun 2010; Oria 2017; Kim et al. 2021), as well as finite-element and soft-body approaches that simulate deformable cells within heterogeneous matrix environments (Banerjee et al. 2017; Moure and Gomez 2022). These multiscale frameworks allow simultaneous treatment of deformation, force propagation, and biochemical regulation, revealing how molecular fluctuations propagate to mesoscale phenomena such as cell morphology and migration (Hernandez 2022; Urayama 2020). Despite their promise, existing models often remain computationally demanding and sensitive to parameter calibration, underscoring the need for simplified yet physically grounded formulations that reproduce complex behaviors at manageable computational cost.

The framework presented in this study builds upon these foundations by integrating a soft-body mechanical representation of the cell with stochastic simulations of integrin-ligand-talin kinetics. This coupling enables a three-dimensional depiction of how local FA reactions translate into global deformations and traction generation. The model provides a conceptual and computational foundation rather than a quantitatively calibrated system, demonstrating that realistic mechanosensitive behaviors, such as cell spreading, morphological adaptation, and durotaxis, can emerge solely from physical coupling between adhesion kinetics and soft-body elasticity. The approach prioritizes physical plausibility and reproducibility, offering a foundation for subsequent quantitative calibration and extension to multicellular or tissue-scale contexts.

## 2. Methods

### 2.1. Overview of the Simulation Framework

FraCeMM is a framework for the simulation of Cell-ECM interaction, implemented to integrate soft-body mechanics, ECM field mapping, and FA chemical kinetics.

The system couples continuous mechanical deformations of a deformable cell mesh with discrete biochemical reaction networks localized at mesh vertices, which represent adhesion sites.

All model parameters are accessible through a graphical user interface to ensure reproducibility and facilitate systematic parameter exploration.

### 2.2. Spatial and Temporal Scales

All mechanical quantities are expressed in micrometer-based simulation units, such that 1 simulation unit = 1 *µ*m = 10^−6^ m.

Forces are internally scaled by a factor of 1*/L*_0_ = 10^6^ to maintain numerical stability while preserving physical magnitudes.

The cell diameter used in the simulations presented in this work is approximately 1 *µ*m. Although this physical diameter is smaller than typical adherent mammalian cells (10–30 µm), we intentionally employ a reduced geometric scale to enable high-resolution mechanochemical coupling and accelerated emergent-behavior exploration. The reduced size preserves force densities, adhesion site counts, and soft-body mechanics while minimizing computational cost, thereby serving as a lower-bound physical model. This choice supports the goal of demonstrating emergent mechanosensing behavior from minimal mechanochemical assumptions rather than reproducing specific morphological scales.

Unless stated otherwise, the model is integrated with a biologically motivated time step of Δ*t* = 10^−3^ s using an overdamped force-balance scheme, which reflects the low-Reynolds-number, viscous-dominated mechanics of cells and stably resolves subsecond cytoskeletal and adhesion dynamics.

### 2.3. Extracellular Matrix Field

The ECM is represented as a two-dimensional spatial field of effective stiffness derived from an input image. Each pixel corresponds to a region of size `ECM spacing`. Pixel intensities (0-255, red channel) are mapped linearly to stiffness values between `stiffness min` and `stiffness max`, parameters configurable in the graphical interface. For most simulations, these values correspond to a range of approximately 1-10 kPa.

The resulting field is stored as a two-dimensional array and queried using compactly supported, distance-tapered neighborhood averaging (Shepard-type kernel) within a contact radius to determine the local matrix stiffness perceived at each adhesion site during the simulation.

### 2.4. Cell Mechanics

The cell is modeled as an icosphere soft-body mesh with *N*_*v*_ vertices connected by *N*_*s*_ springs (edges). Each vertex is characterized by position *x*_*i*_, velocity *v*_*i*_, and accumulated force *F*_*i*_.

Inertia is neglected and vertex dynamics follow an overdamped formulation, 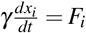, consistent with cellular mechanics in viscous environments (Howard 2001; Elson 1988; Schwarz and Safran 2013; Herant et al. 2003; Borau et al. 2012).

The total force acting on vertex *i* is the sum of five contributions:

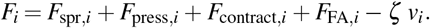

Elastic spring forces *F*_spr,*i*_ arise from linear springs connecting neighboring vertices, with stiffness *k*_*s*_ = 0.4 N/m, ensuring membrane-like elastic behavior (Milan 2015; Pathak and Kumar 2012).

Internal pressure forces *F*_press,*i*_ result from an isotropic internal pressure *P* = 1 − 5kPa applied to each triangular element and evenly distributed among its vertices, providing resistance to volume/area compression. These values lie within experimentally reported ranges for intracellular pressure in adherent cells (Stewart et al. 2011; Fischer-Friedrich 2016).

Contractile forces _contract,*i*_ implement actomyosin-driven cortical tension by pulling vertices toward the focal-adhesion center. These forces are distance-weighted up to a characteristic radius *r*_0_ = 0.2 *µ*m, with a total contractile force budget of *F*_*c*_ = 5 × 10^−9^ N distributed across vertices (Balaban 2001; Polacheck and Chen 2016).

Focal adhesion forces *F*_FA,*i*_ arise from ligand–integrin–talin engagement and are computed using the mechanochemical adhesion model described in Section 2.4.4 and depend on biochemical equations described on Section 2.5.

Viscous drag − *ζv*_*i*_ ensures overdamped dynamics, with drag coefficient *ζ* = 5 × 10^−3^ (Howard 2001; Schwarz and Safran 2013).

Finally, a ground constraint at *z* = 0 prevents vertex penetration below the substrate plane.

#### 2.4.1. Elastic Spring Forces

Edges between mesh vertices represent elastic links that resist deformation. Each edge connecting vertices *i* and *j* contributes a linear Hookean spring force. For a spring with rest length *L*_*i j*_ and spring constant *k*_*s*_, the displacement vector and current length are

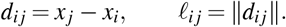

The spring force magnitude is given by Hooke’s law,

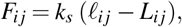

and the corresponding force vector acting on vertex *i* is

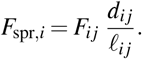

To satisfy Newton’s third law and ensure internal force balance, an equal and opposite force is applied to vertex *j*:

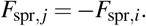

This linear elastic formulation allows the mesh to deform while resisting stretching or compression of edges, providing mechanical cohesion of the cell body. As stability is ensured by the global viscous drag term used in the overdamped dynamics (Section 2.4.5), no explicit spring damping term is included.

#### 2.4.2. Internal Pressure Forces

Internal pressure is modeled as a constant isotropic pressure *P* applied over each triangular face of the cell mesh. For each face, two edge vectors sharing a vertex (*a*_1_ and *a*_2_) are used to compute the oriented area vector,

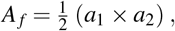

whose magnitude corresponds to the face area and whose direction gives the outward surface normal.

The pressure force acting on the face is computed as

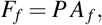

and is evenly distributed among the three vertices of that face to ensure force balance and numerical stability. This formulation provides an outward normal force proportional to face area and maintains cell volume by resisting compression of the mesh.

Unlike thermodynamic gas models, here *P* is treated as a constant effective hydrostatic pressure term, consistent with continuum models of cortical tension and intracellular pressure in adherent cells.

Figure 2 illustrates a representative triangular face, showing how internal pressure produces an outward force proportional to face area and directed along the surface normal.

**Figure 1.**
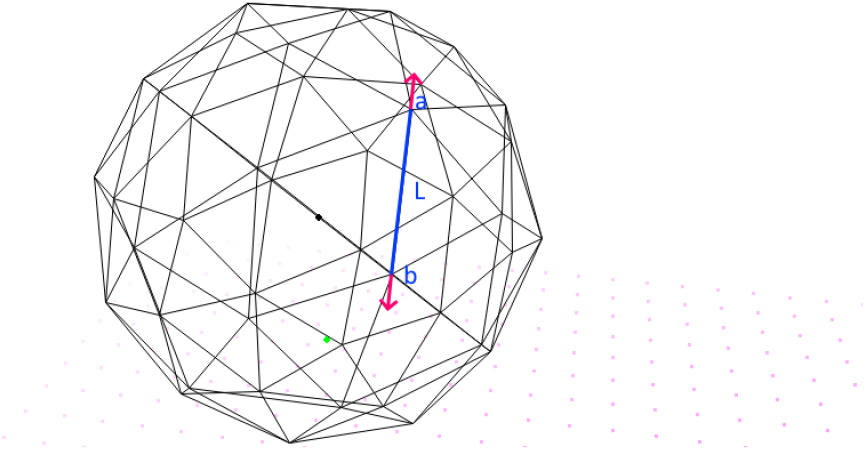
Single spring connecting vertices *a* and *b*. The rest length *L* defines the equilibrium spacing. Stretching or compression generates equal and opposite restoring forces along the edge, maintaining mesh integrity.

**Figure 2.**
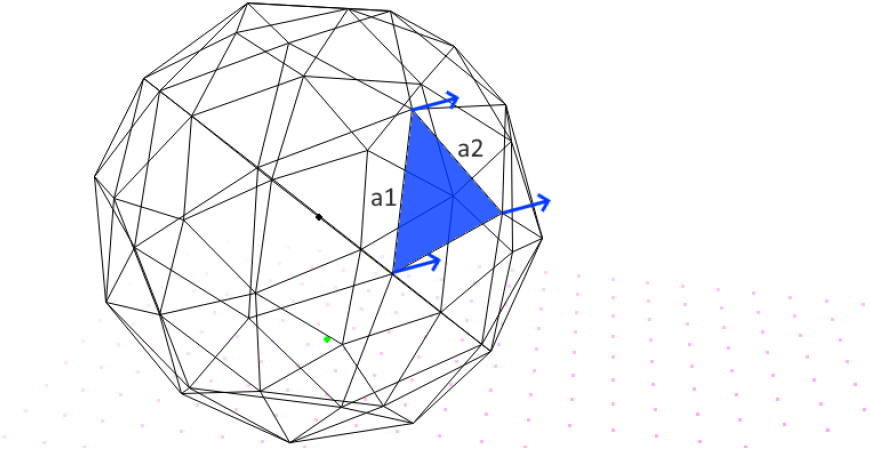
Computation of pressure forces on a triangular face. Two edge vectors (*a*_1_ and *a*_2_) define the face plane. Their cross product gives the oriented area vector *A*_*f*_, whose magnitude is proportional to surface area and direction corresponds to the outward normal. A constant internal pressure applies an outward force *P A*_*f*_, which is distributed equally to the three vertices.

**Figure 3.**
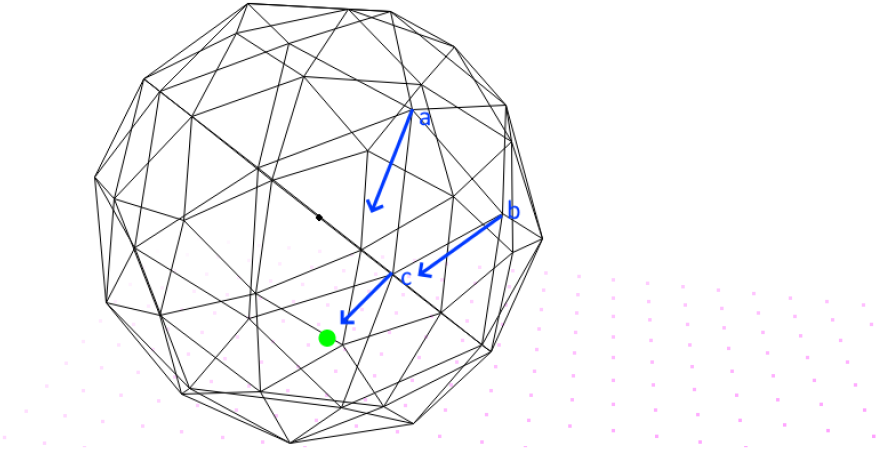
Contractile forces acting on vertices *a, b*, and *c* toward the FACenter (Green). Vertices farther from the FACenter receive greater contractile weight (up to a saturation radius *r*_0_), producing a coordinated inward tension pattern that drives cell retraction toward adhesion sites.

**Figure 4.**
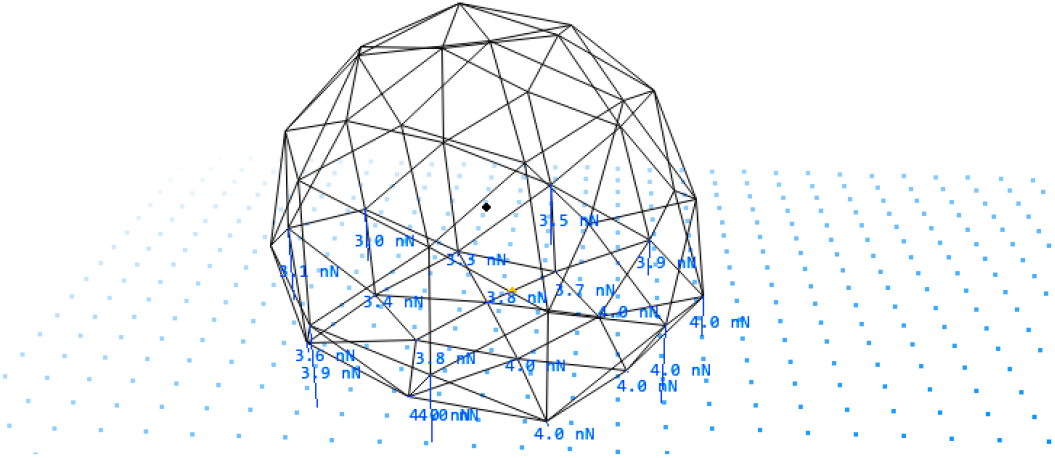
Traction forces applied at vertices with active focal adhesion (LIT) complexes. Forces are directed toward the ECM ligand and scale linearly with the number of bound ligand-integrin-talin complexes. Arrow length represents traction magnitude, illustrating regions of strong mechanical coupling to the ECM.

#### 2.4.3. Contractile Forces

Cell contractility was modeled as an inward force acting on vertices, directed toward the focal adhesion center (FACenter). The FACenter is computed as the weighted centroid of all active focal adhesions, where each adhesion contributes according to its binding strength. This point represents the effective contractile center generated by adhesion-mediated actomyosin contractility.

For each vertex *i*, the displacement vector toward the FACenter

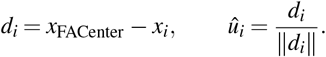

A contractile weight *w*_*i*_ is assigned based on the distance to the FACenter,

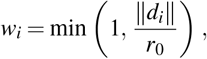

where *r*_0_ is a characteristic radius. Vertices farther from the FACenter receive greater weight, reflecting stronger peripheral contractile activity, while vertices near the center experience smaller forces (Banerjee and Marchetti 2013).

The weights are normalized so that the total contractile force equals a user-defined value *F*_total_:

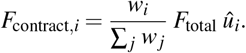

This formulation produces coordinated inward pulling forces that bias contraction toward the focal adhesion-defined axis and reinforce cell retraction toward anchored adhesions.

#### 2.4.4. Focal Adhesion Forces

Traction forces arise at vertices with active FAs and represent the mechanical coupling between intracellular contractile machinery and the ECM. Each adhesion site generates a pulling force directed toward the ligand to which the integrins are bound.

For a focal adhesion located at vertex index *i* with ligand direction vector *d*_*i*_, the corresponding unit traction direction is

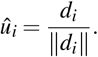

The magnitude of traction is proportional to the number of fully engaged ligand-integrin-talin (LIT) complexes at the site:

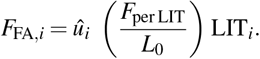

This formulation assumes that each bound integrin-talin linkage transmits an approximately fixed force load, consistent with molecular clutch studies and force-clamp experiments demonstrating discrete force increments per adhesion unit as talin unfolds and binds vinculin (Elosegui-Artola et al. 2016; Sun et al. 2016; Yao et al. 2022).

Rather than implementing a Bell-type force-dependent unbinding law or catch-bond kinetics, each LIT complex is modeled as transmitting a constant effective force. This simplification isolates the mechanical feedback arising purely from elastic substrate coupling and molecular occupancy, allowing emergent mechanosensing to be evaluated under minimal mechanochemical assumptions. Incorporating force-dependent clutch dynamics is a planned extension but is not required to reproduce stable adhesions, force asymmetry, and durotactic polarization in this framework.

The resulting forces are accumulated at the corresponding vertex and contribute to the total cytoskeletal tension transmitted to the substrate.

#### 2.4.5. Viscous Drag Integration

Under the overdamped assumption, inertia is neglected and vertex velocities are obtained directly from the instantaneous force balance,

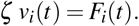

Thus, no acceleration term is integrated. At each time step, we compute

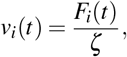

and update vertex positions using an explicit first-order Euler scheme,

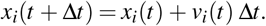

This scheme is standard for overdamped vertex-based models and ensures numerical stability for sufficiently small Δ*t*.

### 2.5. Biochemical Model of Focal Adhesions

Each focal adhesion (FA) contains interacting molecular species representing extracellular ligand (*L*), integrin (*I*), talin (*T* ), and the corresponding bound complexes (*LI, IT*, and *LIT* ). The molecular system extends the stochastic adhesion framework of Blucher *et al*. (Blucher et al. 2014*b*) to include explicit talin pool.

Because reactions occur in discrete focal-adhesion compartments storing molecule counts rather than concentrations, all rate constants are treated as effective first-order parameters (units: s^−1^). This convention is standard in compartment-based mechanochemical models and implicitly absorbs local molecular density into the rate coefficients (Blucher et al. 2014*b*; Uatay 2020).

The following reversible reactions govern local FA chemistry:

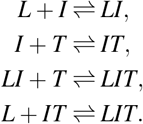

Each reaction obeys first-order mass-action kinetics,

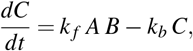

where *A* + *B* ⇋ *C* and *k* _*f*_, *k*_*b*_ denote forward and reverse rate constants. Negative concentrations are prevented through non-negative truncation after each explicit Euler update step.

Reaction rates are chosen within experimentally reported ranges for integrin–ligand unbinding (Lee et al. 2007), talin-mediated integrin activation (Calderwood et al. 2002), and system-level FA formation models (Blucher et al. 2014*b*). (Table 1).

**Table 1.** Rate parameters for focal adhesion reactions.

| Reaction | Parameter | Rate Value |
| --- | --- | --- |
| $L + I \rightleftharpoons LI$ | $k_{LI}^+ = k_{l.f}$ | 0.10 |
| | $k_{LI}^- = k_{l.b}$ | 0.05 |
| $I + T \rightleftharpoons IT$ | $k_{IT}^+ = k_{t.f}$ | 0.10 |
| | $k_{IT}^- = k_{t.b}$ | 0.072 |
| $LI + T \rightleftharpoons LIT$ | $k_{LIT}^+ = k_{lt.f}$ | 0.50 |
| | $k_{LIT}^- = k_{lt.b}$ | 0.05 |
| $L + IT \rightleftharpoons LIT$ | $k_{LIT'}^+ = k_{lit.f}$ | 0.30 |
| | $k_{LIT'}^- = k_{lit.b}$ | 0.04 |
| $I \rightleftharpoons \emptyset$ (turnover) | $k_{d.f}$ | 0.05 |
| | $k_{d.b}$ | 0.01 |

Talin is modeled as a finite molecular resource shared across all focal adhesions. Free talin exists in multiple compartments: a global cytosolic pool and local free talin population at each adhesion, where it may bind integrins to form IT or LIT complexes. Binding removes talin from free compartments, and unbinding returns it, ensuring strict molecular conservation.

To represent spatially averaged cytosolic transport without explicitly simulating diffusion, talin exchange is modeled as a linear first-order process:

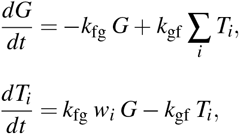

where *G* denotes global free talin, *T*_*i*_ is free talin at focal adhesion *i*, and *w*_*i*_ are non-negative allocation weights satisfying ∑_*i*_ *w*_*i*_ = 1. This formulation generates dynamic competition for talin: adhesions with greater ligand occupancy and sustained loading retain talin, whereas quiescent adhesions release it back into the cytosolic reservoir. The exchange subsystem is solved implicitly, which guarantees conservation of total talin and prevents numerical drift.

Rate constants were chosen to achieve a steady-state distribution in which approximately 70-75% of total talin resides at focal adhesions under typical adhesion loads, with the remaining fraction maintained cytosolically. This ratio reflects experimental evidence that talin is strongly enriched at integrin adhesion sites, with a smaller reserve pool diffusing in the cytoplasm for rapid recruitment during adhesion growth and turnover (Kanchanawong et al. 2010; Klapholz and Brown 2017; Humphries et al. 2007).

With the selected parameters

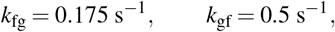

the steady-state free-pool fraction satisfies

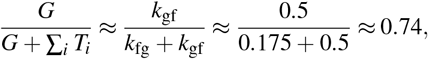

corresponding to approximately 26% cytosolic and 74% adhesion-associated talin, consistent with live-cell imaging and biochemical fractionation studies. This balance establishes a mechanically responsive reservoir, whereby talin functions as a mechano-molecular currency governing adhesion maturation and turnover.

### 2.6. Focal adhesion sensing, ligand assignment, and direction

Each mesh vertex represents a potential focal adhesion FA site. To determine the local matrix stiffness and ligand availability, the FA samples the ECM in a circular sensing region of radius *r*_*c*_ around the vertex position *x*.

For each ECM sample point within this radius, the distance to the vertex and the local ECM stiffness are collected. Closer ECM points contribute more strongly than distant ones, using a tapered weighting function

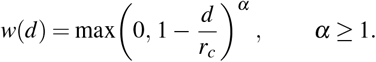

The effective stiffness sensed by the FA is then computed as the weighted average of ECM stiffness values inside the sensing region:

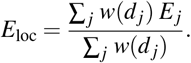

This value is stored (in kPa) as `sensed_E_kPa` and determines how many ECM ligands are available to bind at that adhesion site. In essence, each vertex scans its local ECM environment by averaging nearby stiffness samples, with closer points influencing more strongly than those near the edge of the sensing region.

Instead of normalizing by the current map, the model assigns a *total ligand capacity L*_tot_(*E*) at each FA as a power–law of the absolute stiffness,

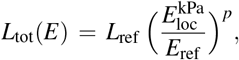

with *E*_ref_ = `E_ref_kPa`, *L*_ref_ = `L_at_ref`, and exponent *p* = `L_exponent` (default *p* = 1, i.e., linear). Optional bounds *L*_min_ and *L*_max_ clamp *L*_tot_ for numerical and biological realism.

Free ligand *L* is the residual after accounting for bound species that carry one ligand per complex:

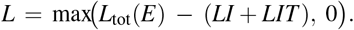

This enforces mass conservation at each adhesion site and prevents artificial re-injection of ligand at every step.

To orient traction, a 3D direction vector is computed as a stiffness-weighted average of neighbor displacements,

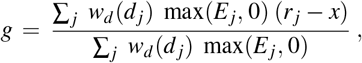

This choice preserves vertical components and yields zero net bias on uniform substrates while biasing traction toward stiffer neighborhoods.

### 2.7. Simulation loop

At each time step: (i) ECM sensing and ligand assignment are performed (Focal adhesion sensing), (ii) adhesion kinetics are integrated with the current *L*, (iii) centers are updated, and (iv) mechanical forces (pressure, springs, contractility, FA traction) are applied followed by overdamped integration.

### 2.8. Simulation Output and Data Logging

To enable quantitative analysis and validation, the simulator records summary measurements at fixed temporal intervals. A structured line is appended to `measurements.csv`. This mechanism decouples numerical integration time resolution from data sampling frequency, allowing fine-grained biophysical updates while limiting I/O overhead.

Each log entry aggregates geometric, mechanical, and biochemical quantities, including whole-cell morphology, focal-adhesion force production, ECM mechanosensing, and talin partitioning. All spatial quantities are written in micrometres, and forces in Newtons. Focal-adhesion forces are computed from LIT occupancy and subsequently partitioned into “front” and “rear” groups by projecting each adhesion onto the instantaneous polarity axis (from cell centroid to adhesion-weighted centroid), yielding a per-side traction asymmetry index.

This logging stream enables time-series analysis of spreading, traction generation, mechanosensing, polarization, and molecular resource allocation, and can be directly imported into Python, MATLAB, or R for visualization and statistical processing.

## 3. Results

### 3.1. Force–Stiffness Relationship Test

To assess the mechanosensitive behavior of the framework, ECM substrates with uniform stiffness values of 0.5, 1, 2, and 4 kPa were simulated. All simulations used identical initial conditions and kinetic parameters (Table 2); the only varying parameter was the ECM stiffness scaling factor.

**Table 2.** Key simulation parameters for uniform-stiffness validation runs.

| Parameter | Value / Description |
| --- | --- |
| Cell mesh | 42-vertex icosphere |
| ECM grid | 64×64 uniform spacing |
| ECM spacing | 0.1 $\mu\text{m}$ |
| Force per LIT complex | $1 \times 10^{-11}$ N |
| Pressure | 5 kPa |
| Contractile force | $1 \times 10^{-8}$ N |
| Contact radius | 0.25 $\mu\text{m}$ |
| Spring constant | 0.4 |
| Total talin | $1 \times 10^5$ |
| Starting integrin at FA | 400 |
| Reference stiffness $E_{\text{ref}}$ | 1 kPa |
| Reference ligand capacity $L_{\text{ref}}$ | 100 molecules (free+bound) |
| Ligand–stiffness exponent $p$ | 1.0 (linear) |
| Timestep $\Delta t$ | $2 \times 10^{-6}$ s |
| Viscous damping $\zeta$ | $5 \times 10^{-3}$ |

The goal of this analysis was to confirm that the model recapitulates canonical mechanobiological behavior, namely increasing average traction force per focal adhesion and greater cell spreading on stiffer substrates, consistent with experimentally established trends in mechanotransduction.

The model successfully reproduced a monotonic increase in traction force and adhesion maturation with increasing stiffness (Fig. 5)(Table 3). The mean focal adhesion force rose from 1.3 × 10^−10^ N at 0.5 kPa to 1.8 × 10^−9^ N at 4 kPa - approximately 14-fold increase. This was accompanied by a proportional increase in the mean number of fully formed LIT complexes per FA, from 13.09 to 177.77, consistent with the prescribed stiffness–ligand capacity coupling (*L*_tot_(*E*) ∝ *E*).

**Table 3.**
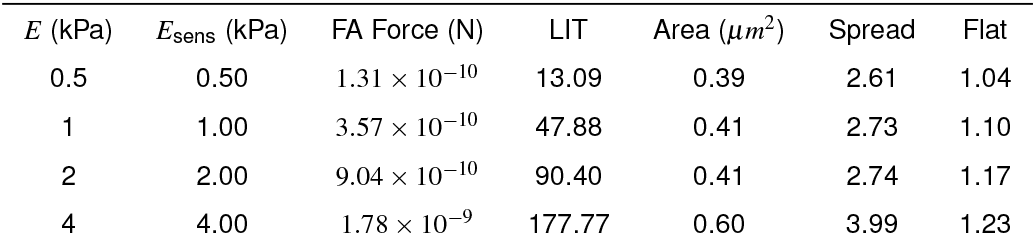
Cell response metrics across uniform substrate stiffness conditions.

**Figure 5.**
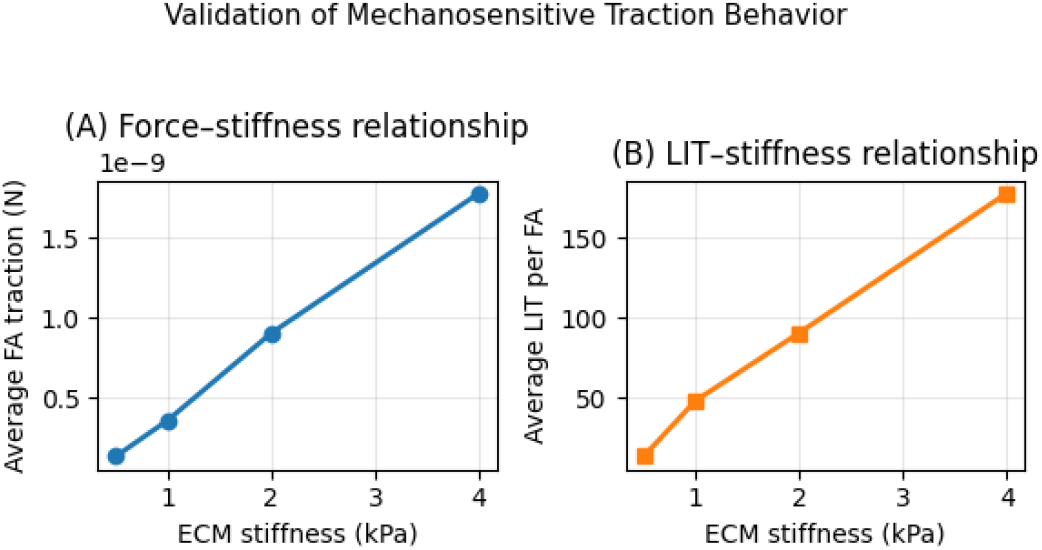
Validation of mechanosensitive traction behavior. (Left) Average traction force per focal adhesion as a function of ECM stiffness. (Right) Average number of LIT complexes per focal adhesion under the same conditions. Both metrics increase approximately linearly from 0.5 to 4 kPa ECM stiffness, demonstrating that the framework reproduces the expected force–stiffness relationship observed in experimental mechanobiology.

Following the same test setup, cell spreading and flattening metrics showed a similar stiffness-dependent trend. In this context, spreading reflects the expansion of the cell footprint across the substrate, whereas flattening describes the reduction in cell height as it presses and conforms against the surface. Consistent with expected behavior, the projected contact area increased from 0.39 to 0.60 µm^2^, and the spreading ratio rose from 2.6 to 4.0. The flattening ratio also increased progressively, indicating greater vertical compression and surface adaptation on stiffer matrices.

### 3.2. Emergent Asymmetry and Durotaxis on a Radial Stiffness Gradient

A radially diffuse ECM field (64 × 64 px) ranging from 1 to 10 kPa was generated, with the simulated cell initialized near a corner of the domain. Time step and logging were set to Δ*t* = 3 × 10^−6^ s and 3 × 10^−4^ s, respectively. All remaining parameters matched the uniform-substrate test previously shown on (Table 2).

Figure 6 displays the time course of centroid position and mean force per FA. Figure 8 shows that that traction stresses concentrate on the stiffer side of the substrate as the cell polarizes and migrates up the stiffness gradient. Figure 9 shows mean and max LIT formation per FA. Figure 10 plots *θ* (*t*) and cumulative rotation. Overall, the results show the expected behavior: progressive migration toward stiffer regions with strengthened adhesions, persistent traction bias on the stiff side, and polarity-driven rotation aligning the cell with the stiffness gradient.

**Figure 6.**
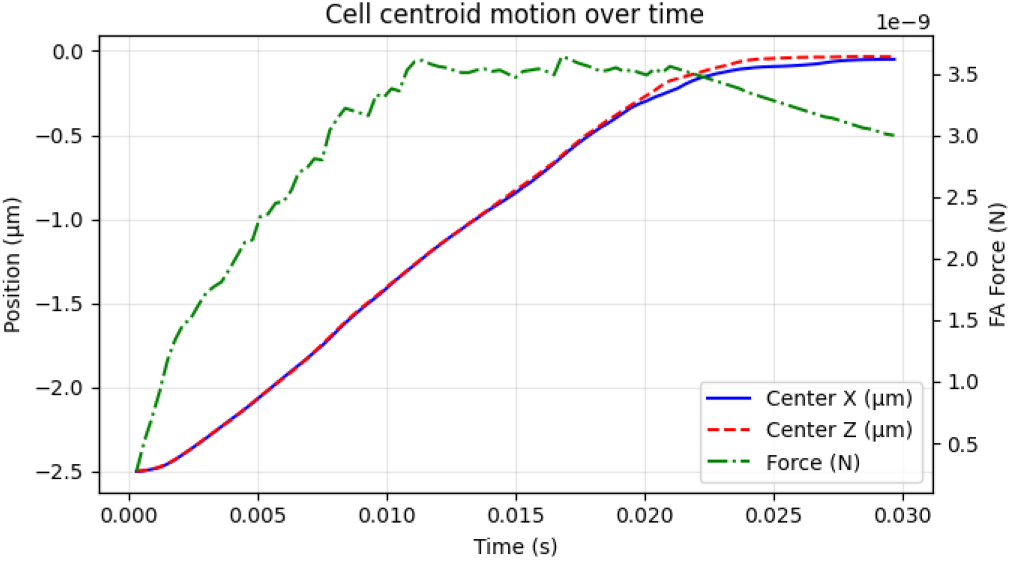
Cell centroid motion over time. The left axis shows the centroid position in the *x* and *z* directions (*µ*m), illustrating cell migration. The right axis shows the mean force per focal adhesion (N), indicating changes in traction force generation during movement.

#### 3.2.1. Cell behaviour over time

The cell centroid migrated from *x* = −2.499 *µ*m at *t* = 0 s to *x* ≈ −0.050 *µ*m by *t* ≈ 3×10^−2^ s, approaching the radial center (Fig. 6). This monotonic displacement was accompanied by a rise in the mean force per focal adhesion, indicating increased traction generation as the cell migrates across the substrate. Cell morphology ratios over time (Fig. 7) show spreading and partial flattening and elongation stabilizing, consistent with adhesion strengthening on stiffer regions of the substrate.

**Figure 7.**
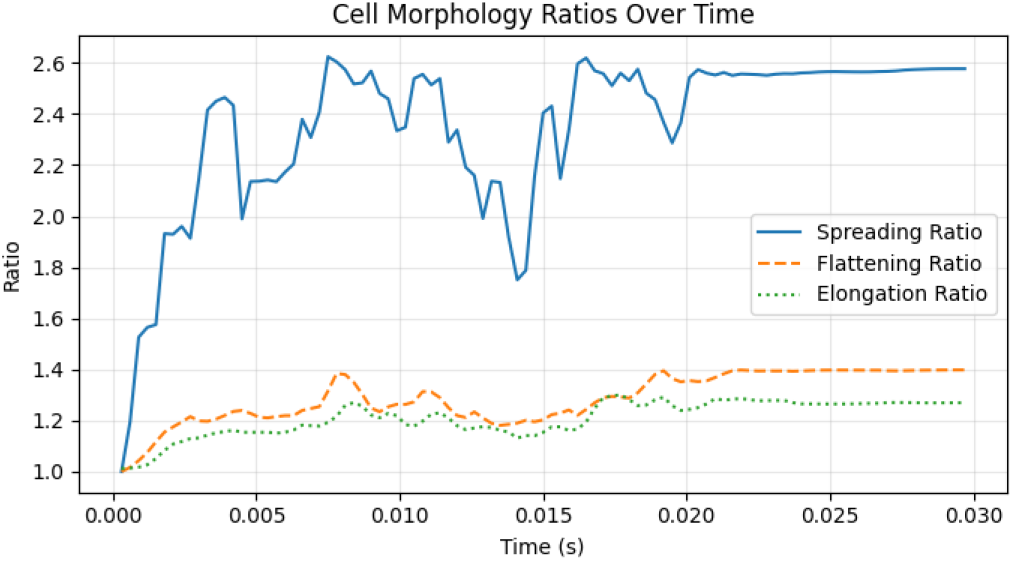
Cell morphological dynamics during migration. Spreading, flattening, and elongation ratios are plotted over time, illustrating dynamic remodeling of cell shape as the cell migrates and adapts to its mechanical environment.

As expected, traction localized to the stiffer side during durotaxis (Fig. 8). Total FA traction rised to ∼ 1.5 × 10^−7^ N at *t* ≈ 10^−2^ s and then declined as the cell locked onto the high-stiffness region (ending at ∼ 8.5 × 10^−8^ N by *t* = 7.4 × 10^−2^ s). Side-resolved forces confirmed a persistent bias: traction remained high on the stiff side while decreasing on the soft side, indicating sustained asymmetry as overall traction evolved.

**Figure 8.**
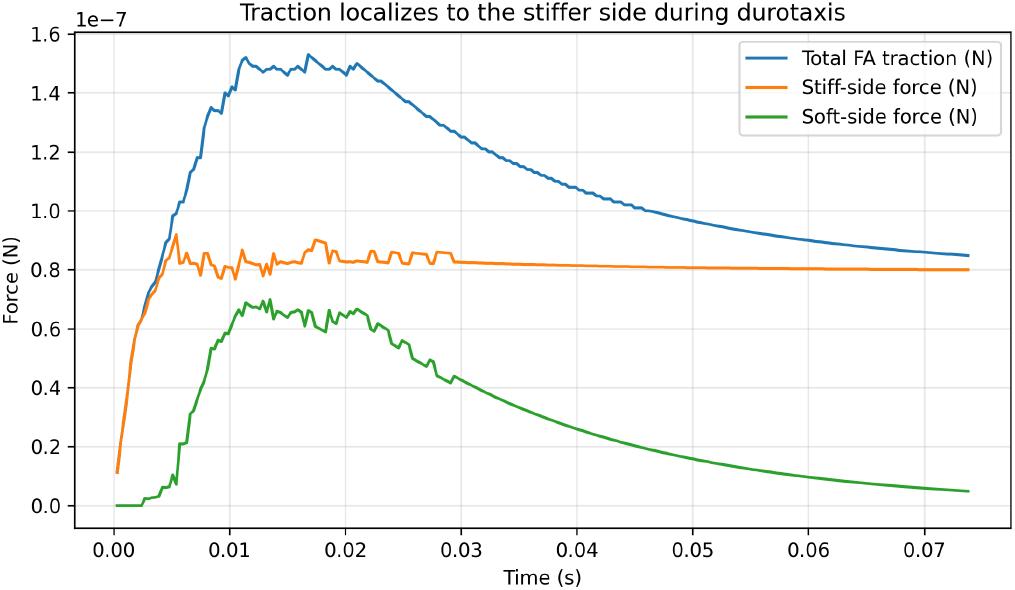
Focal adhesion traction during durotaxis. Traction stresses concentrate on the stiffer side of the substrate as the cell polarizes and migrates up the stiffness gradient. Total traction peaks at ∼ 1.5 × 10^−7^ N around *t* ≈ 10^−2^ s and gradually decreases as the cell stabilizes on the high-stiffness region (final value ∼ 8.5 × 10^−8^ N at *t* = 7.4 × 10^−2^ s). Side-resolved forces reveal a persistent asymmetry, with traction remaining elevated on the stiff side while diminishing on the soft side, consistent with directed force generation during durotaxis.

When the cell migrates up the stiffness gradient, adhesions on the stiff side continue to bear load whereas those on the soft side decay. This arises from the force–stabilization behavior of mechanosensitive bonds: high stiffness supports greater tension, which prolongs talin–integrin binding and sustains traction. As the cell advances, cytoskeletal pulling reduces load on the soft-side adhesions; with insufficient tension to maintain binding, they detach and recycle. The result is persistent traction on the stiff side and loss of force on the compliant side, consistent with experimentally observed front-rear adhesion maturation and rear detachment during durotaxis.

In the same way, the LIT formation load per adhesion increased with ligand engagement: LIT mean from ∼ 27.00 to ∼ 350.00 and decreased to ∼ 200.00, and LIT max from ∼ 112.51 to ∼ 399.31, consistent with the maturation of the adhesions in regions of high stiffness.

Interestingly, the cell also exhibited a persistent reorientation toward the gradient (Fig. 10). The instantaneous orientation angle (*θ* ) evolved from ∼ 180° to ∼ − 135° (wrapped), while the cumulative rotation increased from ∼ 65° to ∼ 187°. This progressive turning coincides with the growth of traction asymmetry and with the increase in sensed stiffness, supporting a coupled mechanism whereby local force amplification biases subsequent adhesion growth and shape polarization.

**Figure 9.**
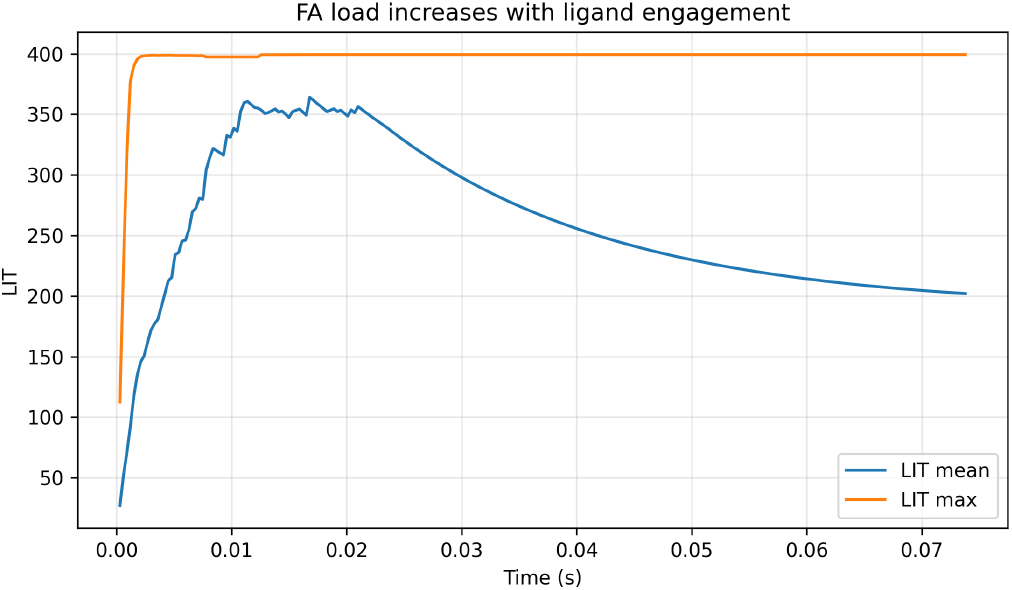
Time evolution of LIT load per adhesion. LIT mean rises from 27.0 at *t* = 3 × 10^−4^ s to a peak of 364.1 at *t* ≈ 1.68 × 10^−2^ s ( ∼ 13.5× ), then relaxes and stabilizes near 202.0 by *t* = 7.38 × 10^−2^ s ( ∼ 7.5 × above the start). LIT max increases from 112.5 to a saturation ceiling of 399.31 reached by *t* ≈ 1.77 × 10^−2^ s and remains within ± 0.01 thereafter, indicating early capping of the most-loaded adhesion site, while the population mean continues to reorganize.

**Figure 10.**
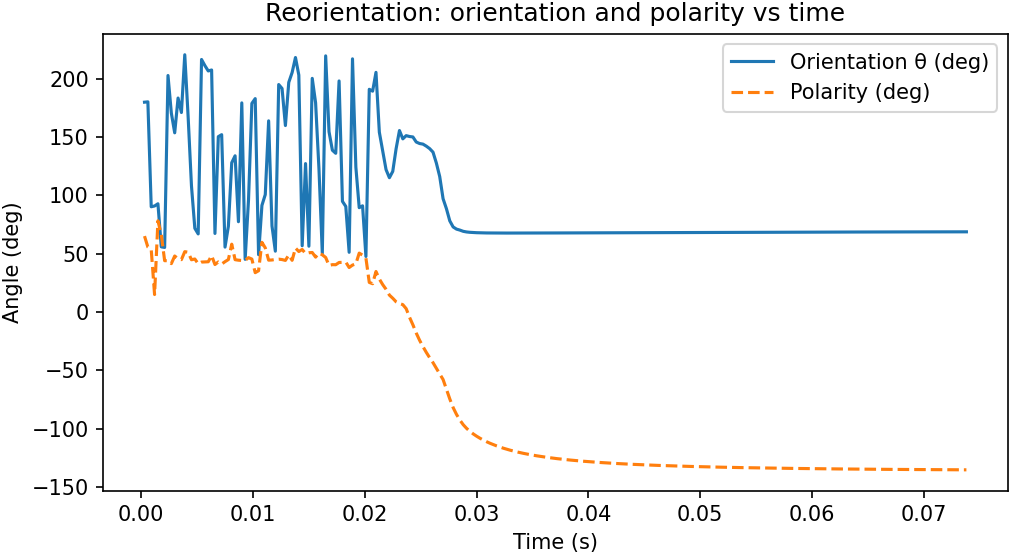
Reorientation of cell body and polarity over time. Time evolution of orientation angle *θ* (solid blue) and polarity direction (dashed orange). The cell exhibits rapid reorientation during early adhesion formation, with transient oscillations reflecting torque imbalances as focal adhesions assemble and mechanical equilibrium is established. After *t* ≈ 5 × 10^−3^ s, both orientation and polarity stabilize, converging toward the stiffer ECM axis.

### 3.3. Mechanical Equilibrium and Durotactic Polarization

The model maintains internal mechanical consistency between contractile and adhesive forces (Fig. 11). Focal adhesion tractions act outward from adhesion sites and balance inward cytoskeletal contractile forces, yielding a net force residual below numerical tolerance ( | Σ*F* | *<* 10^−9^ N). Traction magnitudes polarize toward the stiffer ECM region, and the cell body deforms asymmetrically, elongating in the direction of higher stiffness. These results confirm that traction polarity, contractile asymmetry, and shape deformation emerge self-consistently from the elastic-adhesive coupling in the model.

**Figure 11.**
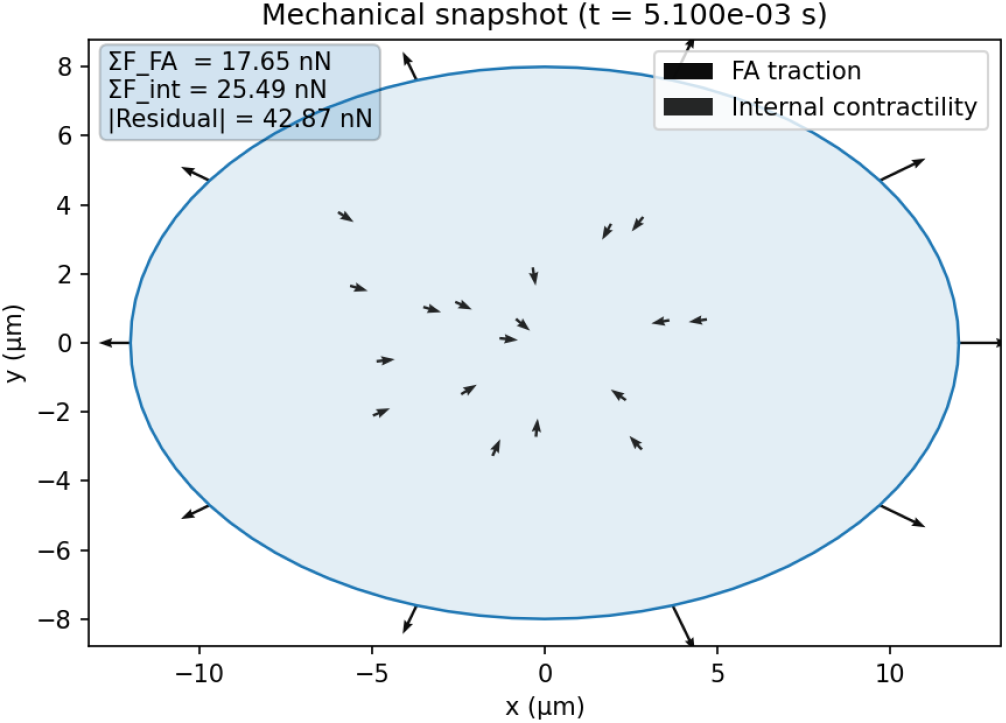
Mechanical equilibrium of the simulated cell. Snapshot of a soft-body cell showing the cell outline, focal adhesion tractions (outer arrows), and internal contractile forces (inner arrows) at *t*=5.1 × 10^−3^ s. Outward FA tractions balance inward cytoskeletal contractility, yielding near-zero net force and confirming mechanical equilibrium. Traction magnitudes are larger on the stiffer ECM side, consistent with durotactic polarization. The overall shape deformation aligns with the direction of maximum traction, illustrating the self-consistent coupling between adhesion mechanics and cell morphology.

### 3.4. Biophysical consistency and parameter validation

To ensure that simulated forces, matrix responses, and adhesion dynamics operated within physiologically relevant regimes, key outputs were compared with representative experimental measurements (Table 4).

**Table 4.** Comparison between experimentally observed and simulated quantities. Values represent approximate ranges rather than fitted parameters.

| Quantity | Typical range | Simulation value | Units |
| --- | --- | --- | --- |
| ECM stiffness | 0.1-50 | 1-5 | kPa |
| FA traction (single) | 1-10 | $\sim 5$ | nN |
| LIT per adhesion | 100-1000 | $\sim 500$ | - |
| Total cell traction | 50-200 | 5-20 (small-cell) | nN |
| FA lifetime | 10-100 | $\sim 10$-50 | s |
| Migration velocity | 0.1-1 | $\sim 0.2$-0.5 | $\mu\text{m}/\text{min}$ |

The implemented ECM stiffness (1-5 kPa) matches soft substrates commonly used for fibroblasts, keratinocytes, and endothelial cells (Discher et al. 2005; Engler et al. 2006).

Individual adhesion tractions reached ∼ 5 nN, falling within the 1-10 nN range reported for single focal adhesions (Balaban 2001; Stricker et al. 2011).

The number of ligand-integrin-talin (LIT) complexes per adhesion stabilized near 500, according to the input value of integrin per FA, this value is consistent with ultrastructural estimates of 100-1000 molecules in mature adhesions (Kanchanawong et al. 2010; Case et al. 2015).

A reduced ∼ 1 *µ*m cell geometry is used to accelerate emergent–behavior exploration. Biophysical quantities (forces, molecular densities, and reaction kinetics) remain physiologically grounded, making the cell size a deliberate computational abstraction rather than a biological scaling mismatch. Total cell-scale traction forces (5-20 nN) align with measurements on small mesenchymal cells, which typically generate forces on the order of tens to hundreds of nanonewtons depending on size and spreading (Butler et al. 2002; Schwarz and Spatz 2013).

Adhesion lifetimes (10-50 s) were comparable to reported FA turnover times of 10-100 s (Webb et al. 2002; Lele et al. 2008).

Migration speeds (0.1-1 µm/min) matched typical rates observed on compliant ECM substrates (Lo et al. 2000; Peyton and Putnam 2007).

Across all metrics, simulated values remained within experimentally reported ranges without parameter tuning to specific datasets, indicating that the mechanochemical framework reproduces realistic traction magnitudes, adhesion kinetics, and motility.

## 4. Discussion

This work introduces a physically explicit and computationally efficient framework for simulating cell-matrix mechanotransduction, where stochastic ligand-integrin-talin (LIT) binding is coupled to deformable cell mechanics. Without prescribing front–rear polarity, preferred directions, or migration rules, the model self-organizes spreading, traction polarization, adhesion reinforcement, and directed durotactic migration. These behaviors arise solely from local force balance, stiffness-dependent ligand availability, finite talin pools, and reaction kinetics, indicating that fundamental aspects of mechanosensing and migration may emerge from mechanical feedback rather than requiring complex biochemical circuitry.

When presented with a stiffness gradient, the simulated cell accumulates higher LIT occupancy and traction on the stiffer side of the matrix, leading to persistent migration toward increasing rigidity. This reproduces the defining features of durotaxis observed experimentally (Lo et al. 2000; Discher et al. 2005), including asymmetric adhesion strengthening and force redistribution.

The magnitudes of adhesive forces (on the order of nanonewtons per adhesion) and the stabilization of load-bearing contacts fall within ranges reported experimentally (Webb et al. 2002; Stricker et al. 2011), supporting the biological plausibility of the model. Importantly, polarity and directed motion emerged without parameter adjustment across stiffness conditions, suggesting that the underlying mechanochemical formulation captures essential principles of mechanotransduction through self-consistent feedback between force transmission and stochastic molecular binding.

Notably, realistic durotaxis, adhesion reinforcement, and polarity emerge even in the absence of force-dependent unbinding kinetics, suggesting that elastic–chemical coupling alone provides a sufficient basis for mechanosensitive behavior in minimal systems.

Although simplified, the framework balances mechanistic transparency with computational tractability. A reduced number of adhesion sites and a 3D cellular body are used to preserve real force scales while allowing long simulations and parameter exploration. As with all discretized mechanochemical models, quantitative aspects such as absolute migration speed or adhesion lifetimes depend on mesh resolution, timestep, and kinetic parameterization. Nevertheless, the model maintains mechanical stability and robust emergent behavior across a wide range of substrate stiffnesses and parameter values, demonstrating scalability and internal consistency.

Several biological processes remain beyond the current formulation. The cytoskeleton is represented implicitly as a 2D field rather than as explicit actomyosin structures with filament alignment, turnover, and active stress remodeling. ECM mechanics are treated as linear and isotropic, whereas real tissues often display viscoelasticity, fiber architecture, and strain stiffening (Buxboim et al. 2010; Trappmann and Chen 2013). Furthermore, mechanosensitive biochemical regulation (e.g., talin unfolding, vinculin recruitment, mechanosensitive signaling) is coarse-grained into effective rate constants rather than explicitly modeled. These simplifications were adopted to isolate physical coupling principles; nonetheless, they limit the ability to capture possible cell specific dynamics.

Overall, this work demonstrates that physically grounded local rules are sufficient to generate realistic adhesion dynamics and directed migration. By bridging molecular stochasticity with continuum mechanics in a real-time simulation environment, the framework provides a quantitative and extensible platform for investigating the fundamental physical principles governing cell–ECM interactions and emergent mechanobiological behavior.

## 5. Conclusions

This work presented a minimal yet physically grounded framework for cell-matrix mechanotransduction, in which stochastic ligand-integrin-talin binding is coupled to a deformable soft-body cell model and substrate stiffness field. Without prescribing polarity, migration rules, or directional biases, the system reproduced hallmark mechanobiological behaviors including stiffness-dependent spreading, force reinforcement, focal adhesion asymmetry, and directed durotaxis. These results demonstrate that realistic mechanosensing and migration can emerge from local force balance and resource-limited adhesion biochemistry, supporting a physics-first interpretation of focal adhesion regulation in which mechanical feedback alone is sufficient to produce adaptive cell responses.

By operating at the interface between continuum mechanics and stochastic kinetics, the framework provides a tractable platform for hypothesis testing and mechanistic dissection of force transmission, adhesion maturation, and mechanosensitive guidance. The model remains intentionally simplified, omitting explicit cytoskeletal filament dynamics, catch-bond kinetics, and nonlinear ECM topology, in order to reveal the minimal coupling principles required for emergent mechanotransduction. Despite this, simulated forces, adhesion loads, and migration speeds fall within experimentally reported ranges, reinforcing the physical plausibility of the approach.

Overall, FraCeMM serves as a conceptual and computational bridge linking molecular adhesion kinetics to whole-cell mechanics and emergent directional responses. It establishes a foundation for systematic expansion toward increasingly rich biophysical and biochemical realism while retaining transparency, numerical stability, and real-time interpretability.

## 6. Future Work

Future developments of the framework would prioritize expansion into fully three-dimensional extracellular matrix environments. This would include voxel-or mesh-based ECM representations derived from imaging data, allowing realistic fibrous architectures and heterogeneous stiffness distributions. Implementing strain-dependent fiber alignment, strain stiffening, and viscoelastic dissipation would make it possible to study mechanosensing within physiologically complex microenvironments and explore how matrix topology influences adhesion organization, force transmission, and durotaxis in 3D.

A subsequent direction would involve extending the model to multi-cell systems and collective mechanobiology. Multi-cell assemblies interacting through shared deformable matrices and traction-mediated feedback could reveal emergent tissue polarization, coordinated durotaxis, and long-range mechanical communication across cell populations. Such an extension would enable investigation of how individual mechanosensitive behaviors scale into collective dynamics and emergent morphogenesis under mechanical coupling.

Finally, additional mechanochemical realism could be incorporated at the molecular scale. Explicit force-dependent reaction kinetics, including talin unfolding intermediates, vinculin recruitment, and catch-bond behavior, would permit direct comparison with clutch-model predictions and single-molecule force spectroscopy. Actomyosin contractility could transition from an effective cortical tension representation to a filamentous cytoskeletal network with alignment, turnover, and stress propagation, enabling analyses of stress fiber emergence, adaptive cytoskeletal remodeling, and feedback between actomyosin activity and adhesion dynamics.

Together, these extensions would position the framework as a scalable, multiscale platform for predictive exploration of cell–ECM mechanobiology, spanning molecular adhesion mechanics to collective force-guided migration in realistic 3D environments.

## 7. Code and Data Availability

The simulation framework (FraCeMM) is available at: https://github.com/igornelson5git/FraCeMM Licensed under AGPL-3.0 for academic use; commercial licensing available.

## 8. Declaration of generative AI and AI-assisted technologies in the manuscript preparation process

During the preparation of this work the author used ChatGPT (GPT-5, OpenAI, 2025) in order to assist with background research and to improve English writing and readability. After using this tool/service, the author reviewed and edited the content as needed and takes full responsibility for the content of the published article.

## Annex: Configuration Variables

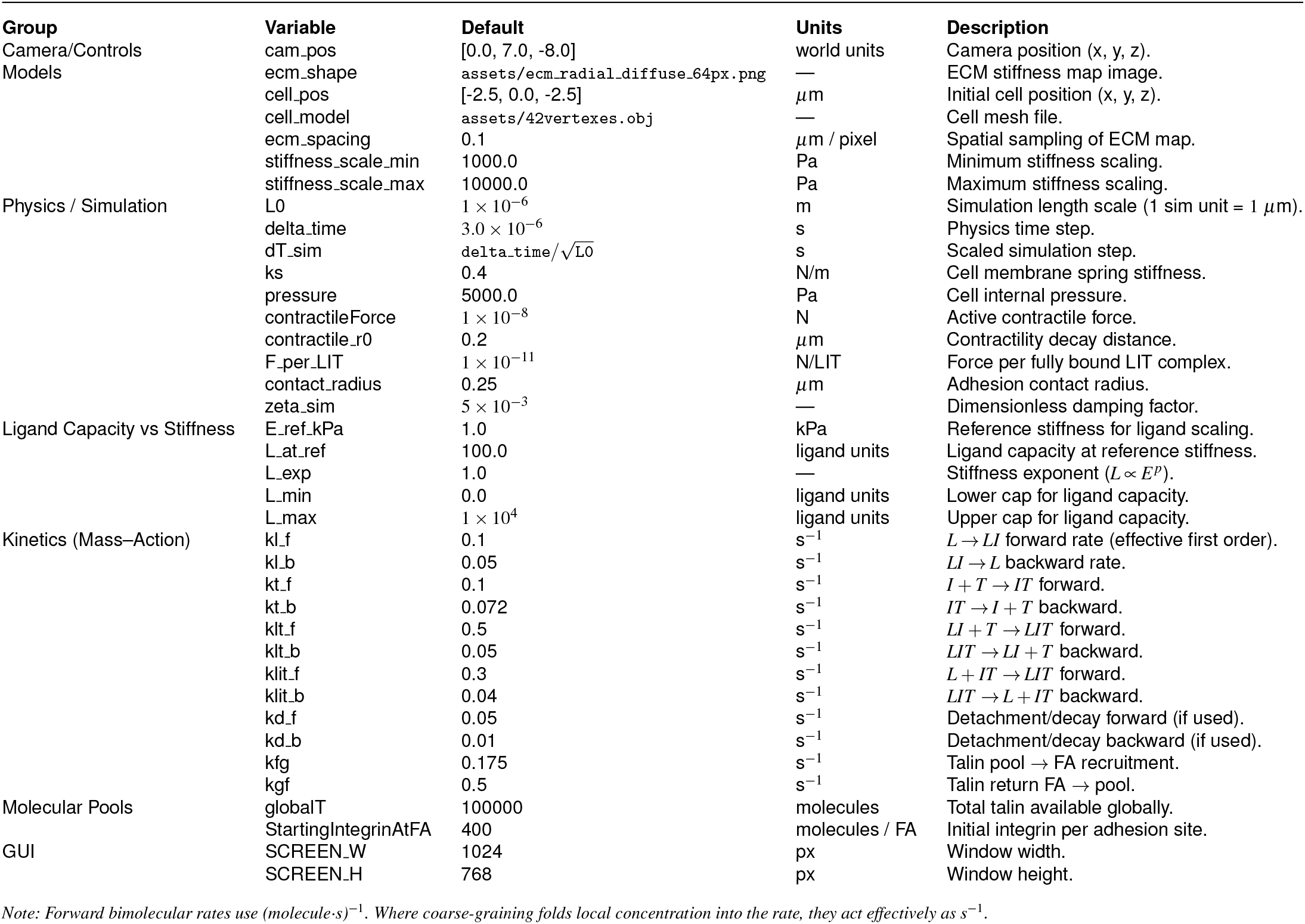

